# PDPN/CLEC-2 axis modulates megakaryocyte subtypes in a hematopoietic stem cell-regulating megakaryocyte-dominant manner

**DOI:** 10.1101/2024.08.14.607891

**Authors:** Rikuto Nara, Hinako Notoh, Tomoyuki Sasaki, Nagaharu Tsukiji, Ayuka Kamata, Nobuaki Suzuki, Atsuo Suzuki, Shuichi Okamoto, Takeshi Kanematsu, Naruko Suzuki, Akira Katsumi, Tetsuhito Kojima, Katsue Suzuki-Inoue, Tadashi Matsushita, Shogo Tamura

## Abstract

**Introduction:** Megakaryocytes are classified into several subtypes including LSP1-positive immune-skewed, MYLK4-positive hematopoietic stem cell (HSC)-regulating, and BMAL1-positive platelet-producing megakaryocytes. Podoplanin (PDPN)-expressing stromal cells generate a microenvironment that promotes megakaryopoiesis in the bone marrow. In this context, PDPN interacts with C-type lectin-like receptor-2 (CLEC-2) on megakaryocyte progenitors, which induces megakaryocyte proliferation. However, the megakaryocyte subtypes developed by the regulation of the PDPN/CLEC-2 axis have not yet been elucidated.

**Materials and Methods:** We established an immortalized bone marrow PDPN-expressing stromal cell line and a PDPN-knockout line (PDPN WT and KO feeder cells, respectively). Bone marrow hematopoietic progenitors were committed to megakaryocytes in co-culture with PDPN WT or KO feeder cells. The number and ploidy of megakaryocytes, resultant platelets, and the polarization of megakaryocyte subtypes were investigated.

**Results:** The number of megakaryocytes was significantly increased in the co-culture with PDPN WT feeder cells compared to that with PDPN KO feeder cells. The megakaryocytes on the PDPN WT and KO feeders showed their main ploidy at 16N∼32N and 8N∼16N, respectively. The number of platelets decreased in the co-culture with the PDPN WT feeder compared to those in the co-culture with the PDPN KO feeder. Megakaryocyte subtypes were immunocytochemically detected in *in vitro* differentiated CD41-positive megakaryocytes. For each megakaryocyte subtype, the percentage of MYLK4-positive megakaryocytes significantly increased and the percentage of BMAL1-positive megakaryocytes significantly decreased when co-cultured with the PDPN WT feeder.

**Conclusion:** The PDPN/CLEC-2 axis modulates megakaryocyte subtype differentiation, with a predominance of HSC-regulating megakaryocytes.

## 1. Introduction

Megakaryocytes are mature hematopoietic cells that differentiate from hematopoietic stem cells (HSCs) and their main function is platelet generation. Recently, megakaryocytes were demonstrated to have several unique functions beyond platelet generation, such as the regulation of HSCs [1-3]. Single-cell RNA sequencing revealed that human and mouse megakaryocytes could be divided into three transcriptomic/functional subtypes: platelet-producing, HSC-regulating, and immunomodulatory (immune-skewed) megakaryocytes [4-7]. Megakaryocyte subtypes are distinguished by specific gene expression: *Lsp1* in immune-skewed megakaryocytes, *Mylk4* in HSC-regulating megakaryocytes, and *Bmal1* (also known as *Arntl*) in platelet-generating megakaryocytes [4].

Podoplanin (PDPN) is a mucin-type transmembrane protein that binds to C-type lectin-like receptor-2 (CLEC-2, formally known as CLEC1B) which is expressed on platelets and megakaryocytes [8-14]. Bone marrow PDPN-expressing stromal cells constitute a perivascular microenvironment that promotes megakaryopoiesis and erythropoiesis in mice [15-17]. During megakaryopoiesis, PDPN binds to CLEC-2 on the megakaryocyte progenitors and induces their proliferative expansion. However, the megakaryocyte subtypes involved in the regulation of the PDPN/CLEC-2 axis have not yet been elucidated.

This study aimed to characterize the megakaryocytes that develop in response to the PDPN/CLEC-2 axis. To our knowledge, this is the first report to demonstrate that bone marrow microenvironmental factors such as PDPN modulate megakaryocyte subtype differentiation. The present findings improve our understanding of how megakaryocytes interact with their environment in the bone marrow.

## 2. Materials and methods

### Mice

C57BL/6NcrSlc mice were purchased from CLEA Japan, Inc. (Tokyo, Japan). They were bred and maintained under standard conditions [12 h light/dark cycle with stable temperature (25 °C) and humidity (60%)]. This study was approved by the Animal Care and Use Committee of the Hokkaido University (22-0087).

### Establishment of immortalized bone marrow podoplanin (PDPN)-expressing stromal cell-line and its PDPN KO-line

Bone marrow stromal cells were isolated as previously described [16]. Briefly, bone marrow cells were harvested from the femurs and tibias of young mice (7–12 weeks old). Flushed bone marrow cells were suspended in cold Iscove’s modified Dulbecco’s medium (IMDM; Wako) containing 10% fetal bovine serum (FBS, Thermo Fisher Scientific) and penicillin-streptomycin (Wako) and seeded into 12-well plates or cell-culture dishes. The culture medium was changed daily for 5–7 d to remove non-adhesive cells until proliferation of adhesive bone marrow stromal cells. To immortalize these stromal cells, the human telomerase reverse transcriptase gene (hTERT) was introduced via retroviral infection. The recombinant virus was prepared by transfecting the pBABE-Puro-hTERT retroviral vector (Cell Biolabs Inc.) into Plat-E retroviral packaging cells (Cell Biolabs Inc.) using Lipofectamine 3000 (Thermo Fisher Scientific). After infecting these stromal cells with the resultant virus-containing culture medium, the immortalized stromal cells were cloned by puromycin selection. Podoplanin expression in cloned stromal cells was evaluated by immunocytochemistry and flow cytometry, as described in the following section. The cloned PDPN-expressing stromal cells were cultured in Dulbecco’s modified Eagle’s medium (DMEM; Wako) containing 10% FBS and penicillin-streptomycin.

The PDPN KO-line was generated by genome editing using the Guide-it CRISPR/Cas9 System (#632601; TaKaRa Bio). The genomic target loci (*Pdpn* exon 1) and guide sequence alignments are shown in Fig 1A. Plasmids were constructed according to the manufacturer’s instructions. The constructed plasmid was transfected into immortalized PDPN-expressing stromal cells by electroporation using Gene Pulser Xcell (Bio-Rad). Transfectants were isolated by limiting the dilution. Genomic DNA extracted from the cloned transfectant was subjected to PCR using KOD FX (Toyobo) and the following primer set: Fw, 5′-TCATCTTTTCACAACCCACAAA-3′; Rv, 5′-CAGGAGACTTAGCCCCATCTAA-3′. The PCR amplicons were analyzed by direct Sanger sequencing using the BigDye Terminator v1.1 Cycle Sequencing Kit and an ABI Prism 310 Genetic Analyzer (Thermo Fisher Scientific).

**Figure 1.**
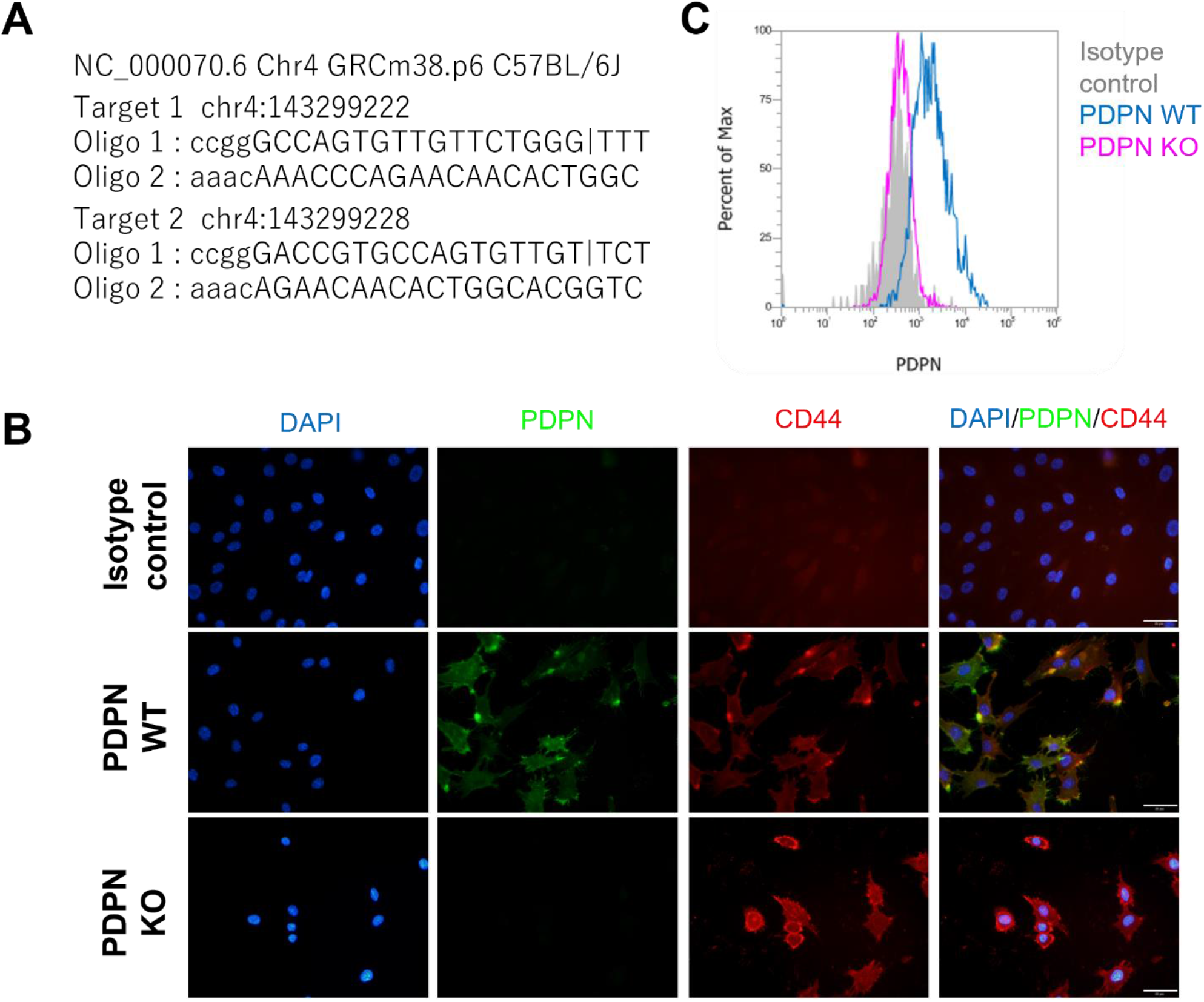
Establishment of immortalized PDPN KO feeder cell line. *A*. Information on genomic target loci and guide sequence alinement in the Guide-it CRISPR/Cas9 System. *B*. Representative immunocytochemistry images of PDPN WT and KO feeders. Blue, DAPI; Green, PDPN; Red, CD44. Scale bar, 100 µm. *C*. Flowcytometry histogram of PDPN expression in PDPN WT and KO feeder cells.

To use each cell line as a feeder, the cells were treated with culture medium containing 10 mg/mL mitomycin C (Nacalai Tesque) for 2 h and collected after washing with phosphate-buffered saline (PBS). The mitomycin C-treated feeders were re-seeded at a density of 0.65 × 10^5^/cm^2^ (2.5 × 10^5^ cells/well in 12-well plates).

### Isolation of bone marrow hematopoietic progenitors

Bone marrow cells were harvested from the femurs and tibias of young mice (8–12 weeks old). The flushed diaphyseal bone marrow was suspended in cold IMDM supplemented with antibiotics and mixed thoroughly by pipetting. After passing the cells through a 40 mm strainer (BD Biosciences), megakaryocyte progenitors were separated by lineage depletion as previously described [16]. Briefly, bone marrow cells were probed with antibodies specific for hematopoietic markers (CD4, CD8, B220, Thy1.2, TER-119, Ly-6G, CD11b, F4/80, antineutrophil antibody, and CD71). Labeled hematopoietic lineage-positive cells were depleted using sheep anti-rat IgG Dynabeads (#11035; Thermo Fisher Scientific). The hematopoietic lineage-negative fraction was harvested as the bone marrow hematopoietic progenitor cells.

### In vitro megakaryocyte differentiation and co-culture with feeder cells

Bone marrow hematopoietic progenitors were cultured in IMDM supplemented with 10% FBS, antibiotics, and 50 ng/mL thrombopoietin (PeproTech) for 5 days to induce megakaryocyte differentiation. On culture day 3, GM6001 (50 mM final concentration; Millipore), a broad-range matrix metalloproteinase inhibitor, was added to the culture medium to suppress the shedding of surface proteins from megakaryocytes. For the co-culturing experiments, hematopoietic progenitors (1.0 × 10^5^ cells per well) were cultured on a feeder cell layer in 12-well plates.

### Immunocytochemistry

The cells were plated and cultured on 15 mm growth cover glasses (Thermo Fisher Scientific) in 12-well plates. The cells were fixed with 4% paraformaldehyde. For the co-culture of megakaryocytes and feeder cells, we carefully pipetted drops of the fixative into the wells and fixed the cells for 1 h to preserve cell-cell contact. The cells were then permeabilized, if necessary, with 0.5% Triton X-100 in PBS for 10 min. After carefully washing with PBS containing 0.05% Treen 20 (PBS-T), cells were blocked with PBS containing 5% bovine serum albumin (BSA) and 2% goat serum, and incubated with antibodies of anti-PDPN (Hamster mAb, 1:200 dilution, clone: 8.1.1; Santa Cruz Biotechnologies), anti-CD44 (Rat mAb, 1:200 dilution, clone: IM7; Thermo Fisher Scientific), anti-CD41 (Rat mAb, 1:500 dilution, clone: MWReg30; Abcam), anti-LSP1 (Rabbit mAb, 1:100 dilution, clone: EPR5997; Abcam), anti-MYLK4 (Rabbit pAb, 1:100 dilution, MBS2005130; MyBioSource), and BMAL1 (Rabbit pAb, 1:100 dilution, ab93806; Abcam). The secondary antibodies used were anti-Syrian hamster Alexa 488 (for PDPN, 1:1000 dilution), anti-rat Alexa 546 (for CD41 and CD44, 1:2000 dilution), and anti-rabbit Alexa 488 (for LSP1, MYLK4, and BMAL1, 1:2000 dilution) conjugates. The cells were mounted using the VECTASHIELD Antifade Mounting Medium containing DAPI (Vector Laboratories) and observed under an inverted fluorescence microscope (IX73; Olympus or ECRIPSE E600; Nikon). The acquired images were quantitatively analyzed using ImageJ 1.46r software (http://rsb.info.nih.gov/ij/).

### Flow cytometry

Cells were probed with fluorescence-conjugated antibodies or their isotype controls as follows: anti-CD41-FITC conjugate (clone: MWReg30; BD Bioscience), anti-CD42b-PE conjugate (clone: Xia.G5; Emfret), and anti-PDPN APC conjugate (clone: 8.1.1; BioLegend). Flow cytometry was performed using a three-laser Attune NxT instrument (Ex: 405/488/637 nm; Thermo Fisher Scientific). For polyploidy analysis, the cultured megakaryocytes were fixed and permeabilized with cold 70% ethanol. After antibody staining with anti-CD41-FITC conjugate and washing with PBS, the DNA content was stained with propidium iodide solution (0.05 µg/mL propidium iodide and 0.25 µg/mL RNase A in PBS) for 30 min in the dark. For the analysis of *in vitro*-generated platelets, platelets were harvested from the culture medium by centrifugation (1,050 × *g* for 10 min) and resuspended in 200 µL PBS containing 5 mM EDTA and 5% BSA. After Fc blocking with anti-CD16/32 antibody for 15 min (clone: 2.4G2; BD Pharmingen), the platelets were washed and stained with anti-CD41-FITC/anti-CD42b-PE. Immunostained platelets were resuspended in 500 µL PBS containing 5 mM EDTA and 5% BSA. The number of platelets in the 200 µL suspension was counted using Attune NxT.

### Statistics

Quantitative data are depicted as mean ± standard distribution of the mean. Representative data from at least three independent experiments are shown. Comparisons between two groups were performed using the unpaired *t*-test. Statistical analyses were performed using GraphPad Prism 5 (GraphPad Software).

## 3 Results and Discussion

### PDPN/CLEC-2 axis promotes megakaryocyte proliferation and polyploidization, but decreases the resultant platelet generation

To investigate megakaryocyte subtype differentiation via the PDPN/CLEC-2 axis, we first established an immortalized bone marrow PDPN-expressing stromal cell line and a PDPN knockout line (PDPN WT and KO feeder cells, respectively, Fig. 1). Bone marrow hematopoietic progenitors were committed to megakaryocytes in co-culture with PDPN WT or KO feeder cells, and the number and ploidy of megakaryocytes were evaluated (Fig. 2A). In addition, the number of resultant platelets in culture with PDPN WT or KO feeder cells were also evaluated. The number of CD41-positive megakaryocytes was significantly increased in the co-culture with PDPN WT feeder cells compared to that with PDPN KO feeder cells (9.43±0.89 megakaryocytes/field in PDPN WT vs 7.09±0.58 megakaryocytes/field in PDPN KO, Fig. 2B and C). Ploidy analysis showed a different pattern of ploidy distribution between megakaryocytes co-cultured with PDPN WT and KO feeder cells (Figs. 2D and E). The megakaryocytes on the PDPN WT and KO feeders showed their main ploidy at 16N∼32N and 8N∼16N, respectively. In a comparison of megakaryocytes cultured with PDPN WT and KO feeder, the percentage of 8N and 32N megakaryocytes were significantly different (8N: 9.49 ± 3.31% in megakaryocytes with PDPN-WT, 29.95 ± 5.37% in megakaryocytes with PDPN-KO, *p* < 0.0001; 32N: 27.83 ± 8.11% in MKs with PDPN-WT, and 8.85 ± 3.73% in megakaryocytes with PDPN-KO, *p* < 0.0001). The number of CD41/CD42b-positive platelets was decreased in the co-culture with PDPN WT feeder, when compared to those in co-culture with PDPN KO feeder (36.20±3.30 platelets in PDPN WT vs 333.33±80.94 platelets in PDPN KO, Fig. 2F and G). The PDPN/CLEC-2 axis induces megakaryocyte proliferation and alters the ploidy pattern of megakaryocytes, but decreases the resultant platelet generation.

**Figure 2.**
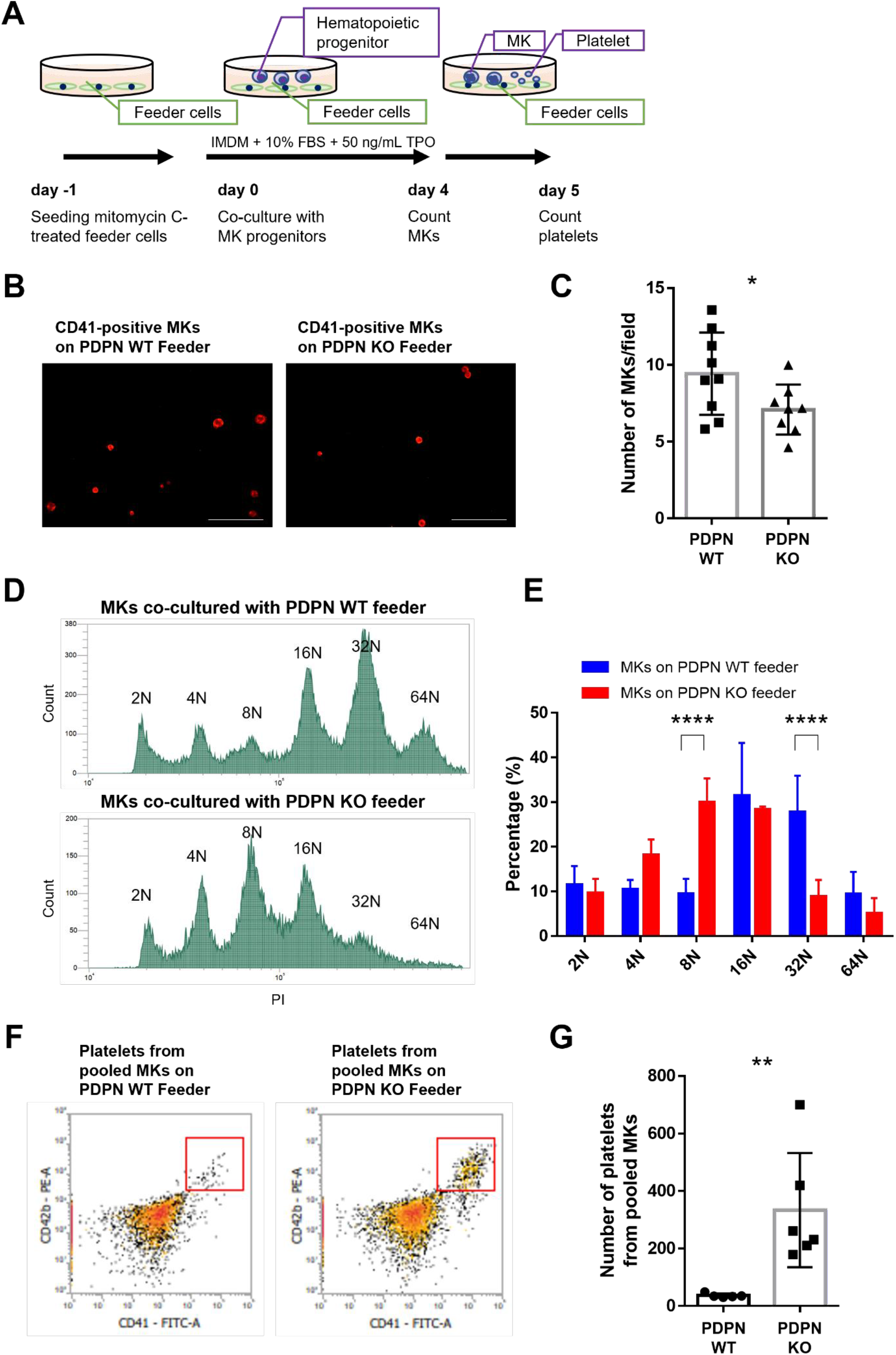
Cellular characteristics of megakaryocytes and their resultant platelets in co-culture with PDPN WT or KO feeder cells. *A*. Graphical illustration of co-culture in bone marrow hematopoietic progenitors with PDPN WT or KO feeder cells. *B and C*. The number of CD41-positive megakaryocytes on PDPN WT or KO feeder cells. Representative immunocytochemical images (B) and their quantitative data (C). Scale bar, 100 µm. *D and E*. Representative ploidy histograms of megakaryocytes co-cultured with PDPN WT or PDPN KO feeder (D and their quantitative data (E). \*\*\*\**p* < 0.0001. n = 3 per group. *F and G*. The number of CD41/CD42b-positive platelets on PDPN WT or KO feeder cells. Representative flowcytometric scatter-grams (F) and their quantitative data (G). *p < 0.05. **p < 0.01. TPO, thrombopoietin; MK, megakaryocyte; PDPN, podoplanin.

### Validation of megakaryocyte subtypes by immunocytochemical detection

Mouse megakaryocyte subtypes present their main ploidy as 2N–8N in LSP1-positive MKs, 16N∼32N in MYLK4-positive MKs, and 8N∼16N in BMAL1-positive MKs [4]. Given these observations, we suspected that the PDPN/CLEC-2 axis is involved in the proliferation of megakaryocyte subtype(s) with nonplatelet-producing characteristics. To investigate whether the PDPN/CLEC-2 axis regulates megakaryocyte subtype differentiation, we immunocytochemically detected each megakaryocyte subtype and assessed its cytological characteristics (Fig. 3). Megakaryocyte subtypes are distinguished by specific marker proteins: LSP1 in immune-skewed, MYLK4 in HSC-regulating, and BMAL1 in platelet-producing megakaryocytes [4]. These subtype marker proteins were immunocytochemically detected in *in-vitro* differentiated CD41-positive megakaryocytes (Fig. 3A). The size of each megakaryocyte subtype was assessed by measuring the major axis diameter (Fig. 3B). The diameter of the LSP1-positive, the MYLK4-positive, and the BMAL1-positive megakaryocytes were 44.80 ± 7.01 µm, 49.29 ± 20.87 µm, and 51.65 ± 24.59 µm, respectively. Immune-skewed LSP1-positive megakaryocytes have a small cytomorphology [4], and megakaryocytes with high ploidy (>16N) and a large cytoplasm exhibit HSC regulation and platelet production functions [18]. In the present study, LSP1-positive megakaryocytes tended to be smaller than the other megakaryocyte subtypes, but the difference was not statistically significant among the megakaryocyte subtypes due to the wide size distribution of MYLK4-positive and BMAL1-positive megakaryocytes. Wang et al. reported that human megakaryocyte subtypes are produced via distinct routes [6]. We consider that the smaller MYLK4- and BMAL1-positive megakaryocytes are cells in the early stages of their differential route. We evaluated the ability of each megakaryocyte subtype to form platelets (Fig. 3C). In the *in vitro* differentiated megakaryocytes, BMAL1 was detected in 83% of the proplatelet-forming megakaryocytes, whereas LSP1 and MYLK4 were not detected in the proplatelet-forming megakaryocytes (Fig. 3D). Based on these observations, we confirmed that the immunocytochemical approach for detecting LSP1, MYLK4, and BMAL1 represented their corresponding megakaryocyte subtypes.

**Figure 3.**
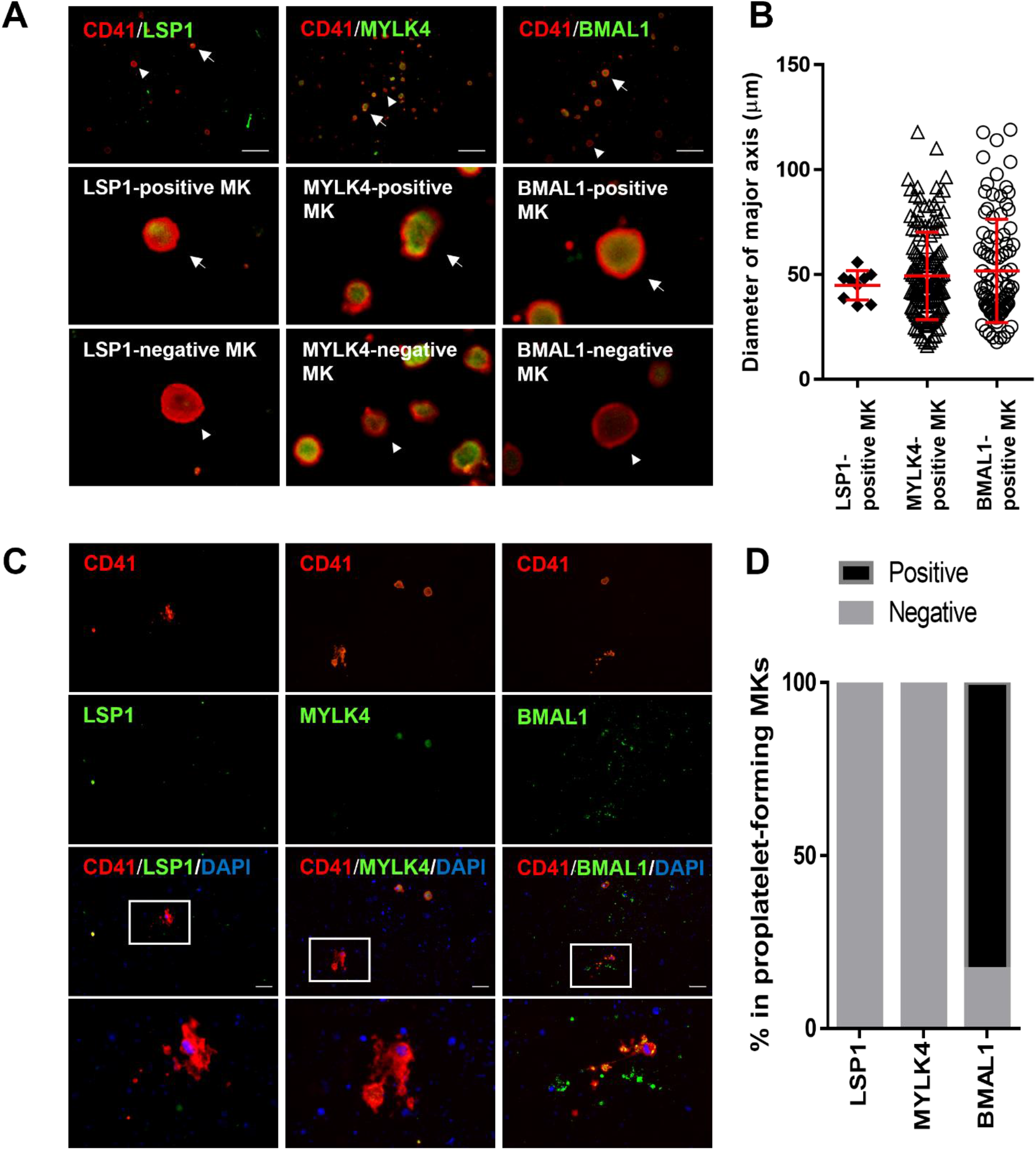
Immunocytochemical detection of megakaryocyte subtypes and their cellular characteristics. *A*. Representative immunocytochemical images of megakaryocyte subtypes with LSP1-positive (immuno-skewed megakaryocytes), MYLK4-positive (HSC-regulating megakaryocytes), and BMAL1-positive (platelet-generating megakaryocytes). Scale bar, 100 µm. *B*. Diameter of major axis in LSP1-, MYLK4-, or BMAL1-positive megakaryocyte. Each megakaryocyte was detected by immunocytochemistry. *C*. Representative immunocytochemical images of proplatelet-forming megakaryocytes. Scale bar, 100 µm. *D*. Frequency of LSP1-positive (3 cells), MYLK4-positive (3 cells), or BMAL1-positive (6 cells) in proplatelet-forming megakaryocytes. Each proplatelet-forming megakaryocyte was detected by immunocytochemistry. MK, megakaryocyte; PDPN, podoplanin.

### PDPN/CLEC-2 axis modulates the balance of megakaryocyte subtypes with a dominance of HSC-regulating megakaryocytes

We evaluated the percentage of each megakaryocyte subtype in co-cultures with PDPN WT or KO feeder cells (Fig. 4). The percentage of LSP1-positive megakaryocytes did not differ between the co-culture with PDPN WT and KO feeders (7.89±0.96% in PDPN WT vs 12.72±2.45% in PDPN KO, Fig. 2B). The percentage of MYLK4-positive megakaryocytes was significantly increased in the co-culture with PDPN WT feeder as compared with that with PDPN KO feeder (67.27±0.32% in PDPN WT vs 57.47±3.09% in PDPN KO, Fig. 2C). In contrast, the percentage of BMAL1-positive megakaryocytes was significantly decreased in the co-culture with PDPN WT feeder as compared with that with PDPN KO feeder (39.99±3.32% in PDPN WT vs 54.90±3.58% in PDPN KO, Fig. 2D). These observations suggest that the PDPN/CLEC-2 axis induces a state of HSC-regulating megakaryocyte predominance in this population.

**Figure 4.**
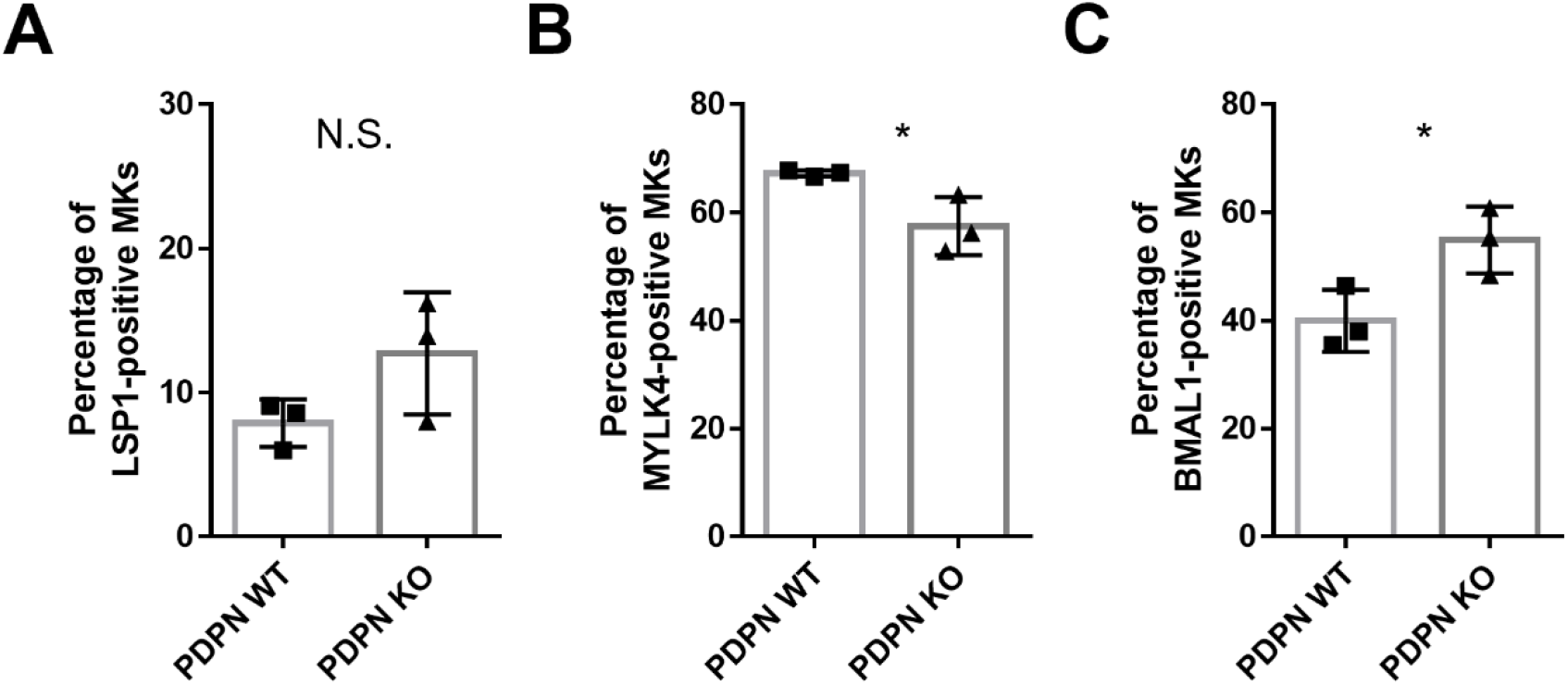
PDPN/CLEC-2 axis modulates the balance of megakaryocyte subtypes with a dominance of HSC-regulating megakaryocytes. *A–C*. Percentages of megakaryocyte subtypes: LSP1-positive (A), MYLK4-positive (B), and BMAL1-positive (C) megakaryocytes. *p < 0.05. N.S., non-significant. MK, megakaryocyte; PDPN, podoplanin.

## 4. Conclusions

This study indicates that the PDPN/CLEC-2 axis modulates megakaryocyte subtype differentiation, with a predominance of HSC-regulating megakaryocytes. These insights suggest that the physiological significance of the bone marrow megakaryopoietic microenvironment comprising PDPN-positive stromal cells is HSC regulation rather than platelet production. Furthermore, the concept of megakaryocyte subtype regulation via exogenous factors suggests the possibility of artificially manipulating megakaryocyte subtype differentiation. One limitation of this study is that we did not elucidate the molecular mechanisms by which PDPN/CLEC-2 regulates megakaryocyte subtype differentiation. The insights gained from this study are mainly based on immunostaining for subtype-specific markers. To further characterize the megakaryocyte subtypes, their gene expression patterns, especially those under the control of the PDPN/CLEC-2 axis, should be investigated. Transcriptome analysis and further cell-biological characterization will lead to a better understanding of megakaryocyte subtype differentiation, providing a new perspective on megakaryocyte biology.

## Abbreviations

CLEC-2: C-type lectin-like receptor-2
HSCs: hematopoietic stem cells
PDPN: podoplanin
PDPN WT: PDPN-expressing stromal cell line
PDPN KO: PDPN knockout line

## Acknowledgments

The authors are grateful to Satoshi Yamasaki, Terumi Nakae, and Hayuki Hayakawa for technical assistance. We would also like to thank Editage (www.editage.jp) for the English language editing.

## CRediT authorship contribution statement

R.N., H.N., T.S., and S.T. Methodology; R.N., H.N., A.K., and S.T., Investigation; R.N., H.N., and S.T. Formal analysis; R.N., H.N and S.T. Writing–original draft; N.T., Nobuaki S., A.S., S.O., T. Kanematsu., Naruko S. A.K., T. Kojima, K.S-I., and T.M. Supervision; S.T. Conceptualization; T. Kojima, T. M., and S. T. Writing–review & editing; R.N., H.N., T.M., and S.T. Data curation; Nobuaki S., A.K., T.M., and S.T. Funding acquisition; S.T. Validation; S.T. Visualization; S.T. Project administration.

## Funding

This study was supported by grants-in-aid provided by the Japanese Ministry of Education, Culture, Sports, Science, and Technology (grant number 22K06881 to S.T. and grant number 22K08518 to A.K.), the National Center for Geriatrics and Gerontology (Research Funding for Longevity Sciences) (grant number 22-10 to A.K.), and JST FOREST Program (grant number JPMJFR2158 to S.T.).

## Declaration of competing interest

The authors declare that they have no conflicts of interest regarding the content of this article.

## Notes

### Competing Interest Statement

The authors have declared no competing interest.

